# Analysis of the lineage of *Phytophthora infestans* isolates using mating type assay, traditional markers, and next generation sequencing technologies

**DOI:** 10.1101/734186

**Authors:** Ramadan A. Arafa, Said M. Kamel, Mohamed T. Rakha, Nour Elden K. Soliman, Olfat M. Moussa, Kenta Shirasawa

## Abstract

*Phytophthora infestans* (Mont.) de Bary, a hemibiotrophic oomycete, has caused severe epidemics of late blight in tomato and potato crops around the world since the Irish Potato Famine in the 1840s. Breeding of late blight resistant cultivars is one of the most effective strategies to overcome this disruptive disease. However, *P. infestans* is able to break down host resistance and acquire resistance to various fungicides, possibly because of the existence of high genetic variability among *P. infestans* isolates via sexual and asexual reproduction. Therefore, to manage this disease, it is important to understand the genetic divergence of *P. infestans* isolates. In this study, we analyzed the genomes of *P. infestans* isolates collected from Egypt and Japan using various molecular approaches including the mating type assay and genotyping simple sequence repeats, mitochondria DNA, and effector genes. We also analyzed genome-wide single nucleotide polymorphisms using double-digest restriction-site associated DNA sequencing and whole genome resequencing (WGRS). The isolates were classified adequately using high-resolution genome-wide approaches. Moreover, these analyses revealed new clusters of *P. infestans* isolates in the Egyptian population. Monitoring the genetic divergence of *P. infestans* isolates as well as breeding of resistant cultivars would facilitate the elimination of the late blight disease.

## Introduction

*Phytophthora infestans* (Mont.) de Bary is a hemibiotrophic oomycete that causes late blight disease in tomato (*Solanum lycopersicum*) and potato (*Solanum tuberosum*). Globally, many epidemics of tomato and potato late blight have been reported since the end of the 19th century, among which the Irish Potato Famine in the 1840s was probably the worst. *P. infestans* is reported to have originated either in South America [1, 2] or in central Mexico [3, 4]. Although numerous studies have been conducted on late blight [5], *P. infestans* remains a major threat to global food security [6]. *P. infestans* is a heterothallic pathogen with two mating types (A1 and A2), which reproduce sexually through oospores [7]. *P. infestans* also exhibits asexual reproduction, with high genetic variability. Since 1980, the A1 mating type was dominant globally, outside the region of its origin [8, 9]. Then, the A2 mating type appeared in many countries [10]. This was followed by the reappearance of the A1 mating type, which replaced A2 and spread in other regions [11]. This repeated appearance and disappearance of mating types might explain why *P. infestans* is able to break down major resistance genes in host plant species and develop resistance against various fungicides [12].

Breeding of late blight resistant cultivars is one of the most effective strategies to overcome this disruptive disease. Resistance genes and/or DNA markers linked to resistance loci are powerful tools for the selection of resistant lines via marker-assisted selection. To the best of our knowledge, genetic loci have been identified in late blight resistant germplasms of wild tomato, *Solanum pimpinellifolium*, in breeding programs [13-15]. More recently, we identified another resistance locus from a susceptible line of cultivated tomato, ‘Castlerock’, which has a minor effect on the resistance phenotype (Arafa et al. under submission). These results suggest that plants harbor many resistance genes and employ several defense mechanisms against the late blight pathogen. Additionally, the pathogenicity of *P. infestans* is complicated and has evolved over time, probably because of sequence variations in virulence genes, also known as effector genes, which encode proteins that suppress the plant immune system and alter the host response to accelerate the infection process. Therefore, knowledge of the genetic divergence of *P. infestans* isolates is important to control the late blight disease.

Over the last several decades, PCR assays for assessing nucleotide sequence variations in mitochondrial DNA (mtDNA), simple sequence repeats (SSRs; also known as microsatellites), and effector genes have been employed to investigate the genetic diversity of *P. infestans* populations [16-20]. Advances in sequencing techniques, based on next generation sequencing (NGS) technology, have enabled the analysis of nucleotide sequence variations on a genome-wide scale. Since the recent release of the *P. infestans* reference genome sequence [21], reduced-representation sequencing approaches, such as genotyping by sequencing [22] and restriction-site associated DNA sequencing [23], have been used to scan genome-wide sequence variations. Moreover, because of the contiguous decrease in the cost of NGS and increase in its throughput, WGRS analysis has enabled the elucidation of the diversity of *P. infestans* populations on a genome-wide level. The genome-wide genotyping methods have been applied to *P. infestans* populations to reveal their phylogenetic relationship [17, 19]; however, the number of *P. infestans* isolates tested has been limited and insufficient to cover the *P. infestans* populations spread across the world.

A decision support system in agricultural domains would be essential for successful plant disease management. Knowledge of the evolution of *P. infestans* clonal linages, based on DNA analysis, epidemiology, and population dynamics of this destructive plant pathogen [19], would be quite useful for the system. Therefore, in this study, we aimed to characterize *P. infestans* populations collected from Egypt and Japan; neither of these regions has been represented in the genome-wide genotyping analysis of *P. infestans* isolates to date. We investigated the mating types, mtDNA haplotypes, and SSR marker genotypes of the isolates as well as genome-wide genotypes, based on double-digest restriction-site associated DNA sequencing (ddRAD-Seq) and WGRS approaches, to gain insights into the diversity of *P. infestans* populations at the genome-wide level.

## Results

### Mating types in the tested isolates

A total of 80 *P. infestans* isolates were used in this study; these included 62 isolates collected from seven counties in Egypt, and 18 isolates obtained from the National Institute of Agrobiological Sciences (NIAS) in Japan, which were collected from three prefectures (Supplementary Table S1). Among these 80 isolates, 67 were of the A1 mating type, 12 were of the A2 mating type, and one was self-fertile (SF). All three mating types were identified among the 18 isolates obtained from the NIAS, whereas only the A1 mating type was observed among the 62 isolates collected from Egypt (Figure 1 and Supplementary Table S1). In the NIAS collection, the A1 mating type isolates were collected from potato leaves in Hokkaido prefecture in 2013, while the A2 mating type isolates, except four, were collected from tomato leaves and fruits in Ibaraki prefecture in 1991–1992. The remaining three A2 mating type were collected from potato in Hokkaido and from tomato in Sizuoka prefecture. The SF was isolated from tomato in Ibaraki.

**Figure 1.**
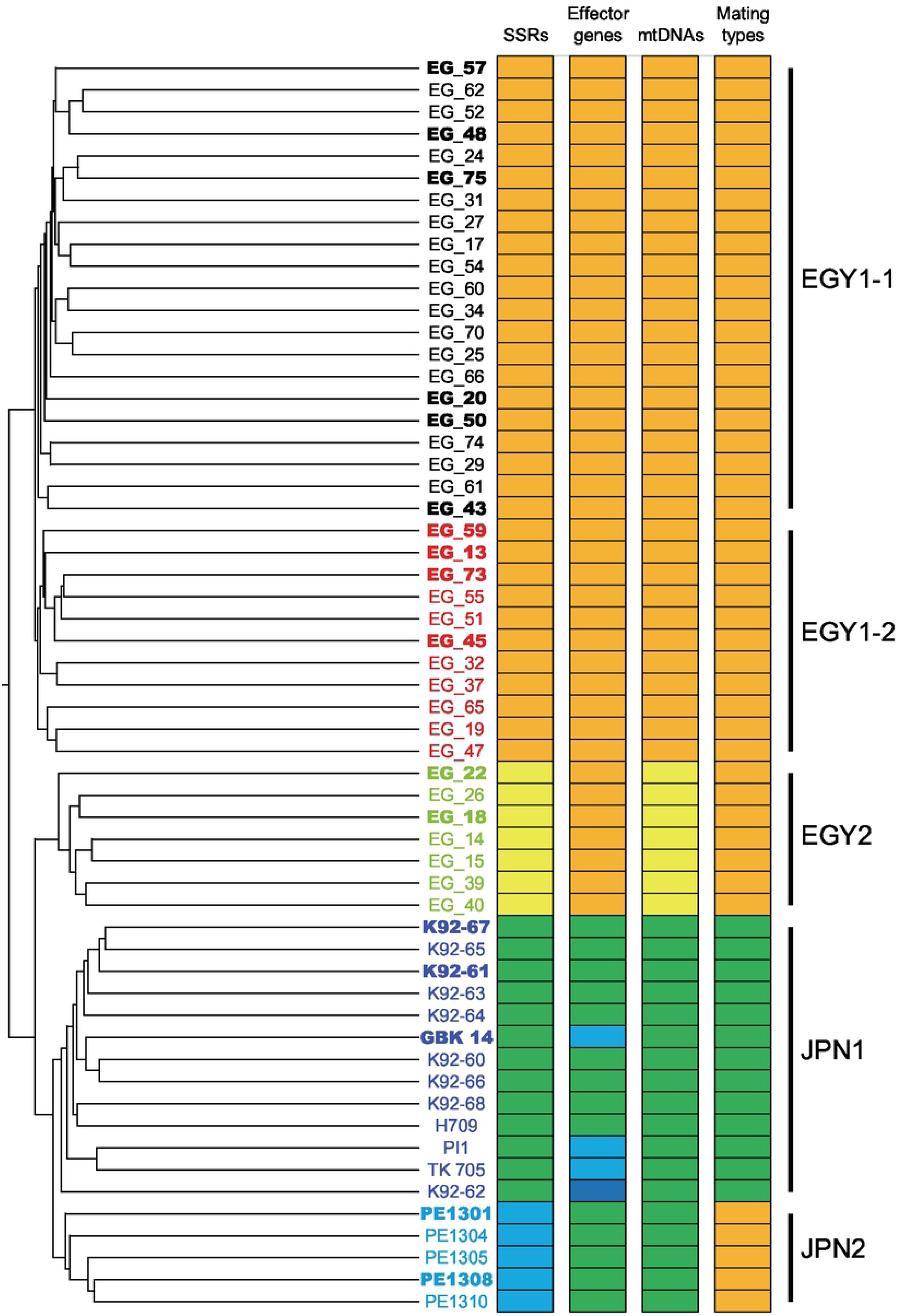
Clustering analysis of *Phytophthora infestans* isolates collected from Egypt and Japan using double-digest restriction-site associated DNA sequencing (ddRAD-Seq) data. Clusters (EGY1-1, EGY1-2, EGY2, JPN1, and JPN2) are indicated with solid bars. Clusters determined using simple sequence repeats (SSRs), effector genes, mitochondrial DNA (mtDNA) polymorphisms, and mating types are shown as colored charts; same colors indicate the members of the same clusters. Isolates indicated in bold were used to perform whole genome resequencing (WGRS) analysis.

### Analysis of mtDNA polymorphisms

The 80 isolates were analyzed by PCR using four restriction fragment length polymorphism (RFLP) markers to detect sequence variations in P1, P2, P3, and P4 regions of the mitochondrial genome (Figure 1 and Supplementary Table S1). The mtDNA polymorphisms classified the isolates into two groups, Ia and IIa. Isolates in each group showed a strong association with their area of collection; isolates in group Ia were collected from Egypt, while those in group IIa were obtained from Japan. Additionally, polymorphisms in two hypervariable regions (HVRs) in the mitochondrial genome were analyzed, which divided the isolates into five groups. The IR1 type was the most dominant among Egyptian isolates (61/62), while three IIR2 and fourteen IIR3 types were observed among the 18 Japanese isolates. The exceptions were one IR2 type in Egyptian isolates (EG_11) and one R1 type in Japanese isolates (K92-62).

### Analysis of effector gene polymorphism

We tested the PCR assay to detect presence–absence polymorphism in five effector genes, *AVR1, AVR2, AVR2-like, AVR3a*, and *AVR4*, across all 80 isolates. DNA amplicons were obtained for four genes (*AVR2, AVR2-like, AVR3a*, and *AVR4*) (Figure 1 and Supplementary Table S1) and a positive control (*NitRed*), but not for *AVR1*. The *AVR4* and *NitRed* genes were amplified from all 80 isolates. However, the remaining three genes showed present/absent polymorphism; *AVR3a, AVR2*, and *AVR2-like* were not detected in 4, 12, and 19 isolates, respectively. All of these three genes (*AVR3a, AVR2*, and *AVR2-like*) were absent in the Egyptian isolate EG_36, while *AVR2-like* was absent in all of the Japanese isolates.

### SSR genotyping analysis

We tested 16 SSR (or microsatellite) markers in the 80 isolates. A total of 54 peaks were detected, ranging from two peaks in five markers (Pi33, Pi70, Pi89, SSR2, and SSR8) to seven peaks in SSR4 (Supplementary Table S1). The 62 Egyptian and 18 Japanese isolates showed 45 and 38 peaks, respectively, for the 16 SSR markers. Clustering analysis divided the 80 isolates into four clusters (Figure 1 and Supplementary Table S1). Two clusters, EGY1 and EGY2, contained 50 and 12 Egyptian isolates, respectively. All of the isolates in the EGY2 cluster were collected in the 2014–2015 growing season, while those in the EGY1 were sampled in 2014–2015 and 2015–2016. The EGY1 and EGY2 clusters showed no relationship with the geographic location, host species, and infected organs, and all isolates in the EGY2 cluster lacked the *AVR2* amplicon. The other two clusters contained only Japanese isolates, with 13 isolates in JPN1 and five in JPN2. Isolates in the JPN1 cluster were collected from 1987–1992, while those in the JPN2 cluster were collected in 2013.

### Discovery of genome-wide single nucleotide polymorphisms (SNPs) using ddRAD-Seq

The ddRAD-Seq approach was used to analyze genome-wide SNPs. A total of 23 Egyptian isolates were removed from subsequent analysis because of low-quality data, probably because the DNA available for analysis was of poor quality and insufficient quantity. An average of 430,489 high-quality ddRAD-Seq reads were obtained per sample from 57 isolates (Supplementary Table S3). Reads were mapped onto the reference genome sequence of the *P. infestans* strain T30-4 (version ASM14294v1) at an average mapping rate of 92.4%, and 996 loci were detected as high-confidence SNPs based on the read alignments.

Clustering analysis, based on SNPs, grouped the tested isolates into four clusters, EGY1, EGY2, JPN1, and JPN2, containing 32, 7, 13, and 5 isolates, respectively, in accordance with the results of the SSR marker analysis (Figure 1). Furthermore, the EGY1 cluster was divided into two sub-clusters, EGY1-1 and EGY1-2, containing 21 and 11 isolates, respectively. Principal component analysis (PCA) supported this result, and PC1, PC2, and PC3 axes explained 11.2%, 4.2%, and 3.8% of the variance, respectively (Figure 2).

**Figure 2.**
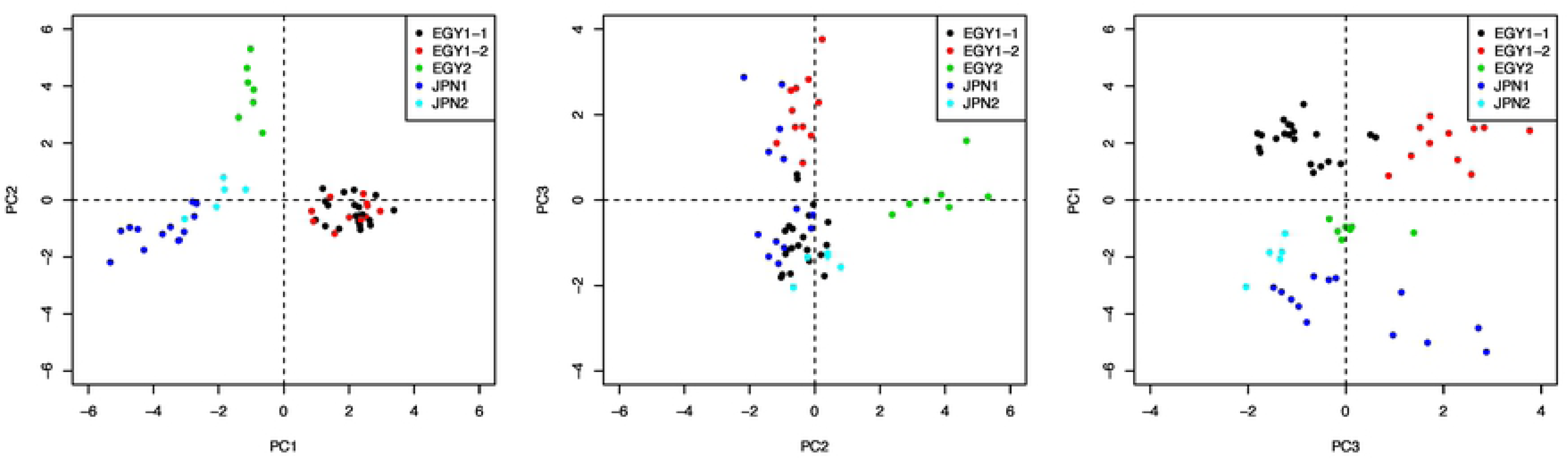
Principal component analysis (PCA) of *P. infestans* isolates. Different colors indicate the five clusters, EGY1-1, EGY1-2, EGY2, JPN1, and JPN2, identified by clustering analysis.

### Whole genome SNP and copy number variation (CNV) analysis

Seventeen isolates representative of each cluster, based on ddRAD-Seq data and sampling location, host, and year, were selected for WGRS analysis. An average of 7.9 Gb sequence data per sample (34.6× coverage) was obtained (Supplementary Table S4). The reads were mapped onto the reference genome sequence of *P. infestans* at an average mapping rate of 94.5%, and 760,928 high-confidence SNPs, including 479,784 transitions and 281,144 transversions (Ts/Tv ratio = 1.7), were detected. The Egyptian isolates EG_18 and EG_73 contained the lowest and highest number of SNPs, respectively. The average density of SNPs was 3.3 per kb. Analysis of SNP annotations using SnpEff indicated that intergenic variants were the most common (606,900; 79.8%), followed by synonymous variants (57,024; 7.4%) and missense variants (56,758; 7.4%) (Supplemental Table S5). A total of 1,267 (0.6%) high- and 56,758 (7.4%) moderate-impact SNPs were identified in 1,085 and 11,289 genes, respectively.

Among the Egyptian isolates, CNVs were detected between EG_18, as a reference, and each of the remaining 16 isolates. Between any two isolates, an average of 5,402 CNVs were detected in each isolate, ranging from 533 CNVs in EG_22 to 6,735 CNVs in K92-61. The average size of a CNV was 4,068 bp, with the longest of 79 kb in EG_75.

To determine the lineage of *P. infestans* isolates, we used the SNP data and data publicly available on 11 isolates [19]. On the basis of 90,7671 SNPs detected in 28 isolates including the 17 isolates from this study and 11 from the publica data, four clusters were generated (Figure 3). Two Egyptian isolates belonging to the EGY2 cluster, EG_18 and EG_22, were grouped with 06_3928A, an isolate collected from England. Two clusters included Japanese and Egyptian isolates belonging to JPN1, JPN2, and EGY1 clusters. The fourth cluster consisted of ten isolates, all of which have been reported previously as members of US-1 and HERB-1.

**Figure 3.**
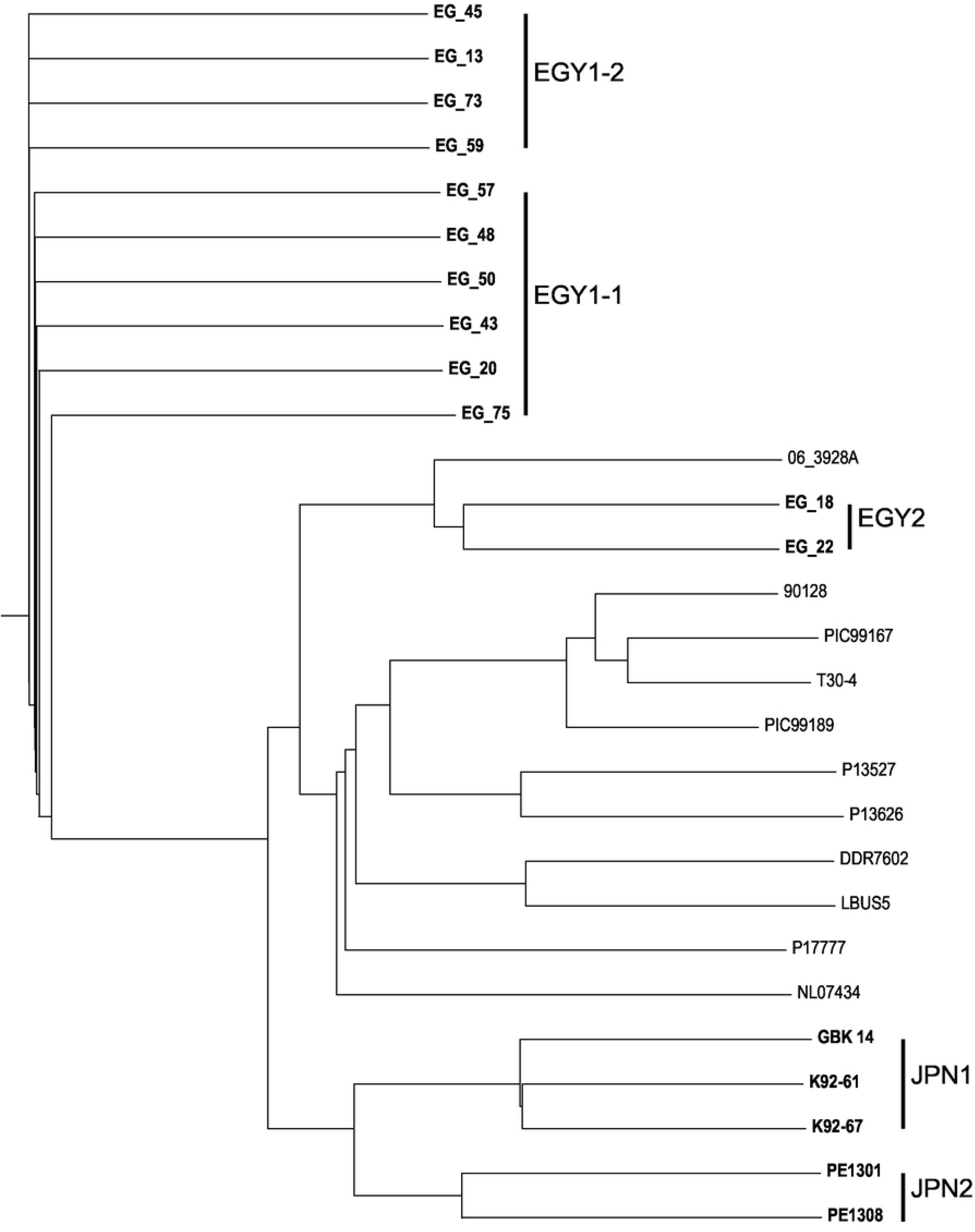
WGRS-based clusters of *P. infestans* isolates. *P. infestans* isolates sequenced in this study are indicated in bold. The remaining isolates represent those whose genome sequences were publicly available.

### Polymorphisms in effector genes

In the *P. infestans* T30-4 reference genome sequence, 594 and 194 genes have been annotated as encoding RxLR- and Crinkler-type effector proteins, respectively. Among these, four genes were identified in the *P. infestans* T30-4 genome sequence: *Avr1* (PITG_16663) in the contig sequence of supercont1.51, *Avr2* (PITG_08943 and PITG_22870) on supercont1.16, *Avr3a* (PITG_14371) on supercont1.34, and *Avr4* (PITG_07387) on supercont1.11. A total of six non-synonymous SNPs causing missense mutations were identified. Four of the six missense mutations caused amino acid substitutions at positions 19, 80, 103, and 124 of PITG_14371 (Avr3a). While the first three mutations were observed in the ten isolates belonging to clusters EGY1-1 and EGY1-2, the fourth mutation was identified in all 12 isolates belonging to the EGY2 cluster, in addition to the isolates in EGY1-1 and EGY1-2 clusters. Among the two remaining missense mutations, one was detected at amino acid position 31 of PITG_08943 (Avr2) in the two isolates belonging to the JPN2 cluster and one isolate (GBK 14) belonging to JPN1, and the other was detected at amino acid position 139 of PITG_07387 (Avr4) in seven isolates belonging to clusters EGY2, JPN1, and JPN2. No SNPs were detected in PITG_16663 (Avr1).

CNVs were detected in *Avr1, Avr2, Avr3a*, and *Avr4*. CNVs of *Avr1* and *Avr4* were observed in 12 isolates of EGY1-1, EGY1-2, and JPN2, while CNVs of *Avr2* were detected in three isolates belonging to the cluster JPN1 and 12 isolates belonging to EGY1-1, EGY1-2, and JPN2. The CNV of *Avr3a* was identified in only one isolate, EG_75, belonging to the sub-cluster EGY1-1.

## Discussion

In this study, we analyzed a panel of 80 *P. infestans* isolates collected from Egypt and Japan using a series of thorough genotyping assays at different levels, including the target gene level (SSRs and RFLPs), genome-wide level (ddRAD-Seq and WGRS), and reproduction biology level (mating type assay). While A1 and A2 mating types were observed among Japanese isolates, the A1 mating type was predominant among Egyptian isolates. This suggests that the A1 mating type is adapted to the Egyptian environment [24], resulting in the absence of A2 mating types needed for sexual reproduction to produce oospores. Nevertheless, Egyptian isolates are more divergent than the Japanese isolates [24]. The unstable nature of *P. infestans* genome may be one of the possible drivers of genetic diversity [21]. In this study, we observed SNPs and CNVs in the genomes of isolates. Some genomic variations may alter the host specificity and geographical location of isolates [17, 25]. According to a recent report [26], CNVs are associated with the emergence of super-fit clones of *P. infestans*. However, effector genes do not diverge among the Egyptian isolates, possibly because most of the tomato and potato cultivars in Egypt are susceptible to late blight, leading to minimal selection pressure and low mutation rates. Nonetheless, the population structure of *P. infestans* isolates, regardless of their reproduction mode, was broader than that suggested by previous studies on American and European isolates (Figure 1). This finding was supported by our WGRS data (Figure 3), which was able to directly compare the isolates from other studies [19, 27-29]. The ddRAD-Seq analysis also provides information on population structure [30-32]; however, the ability to compare data depends on the choice of restriction enzymes used for library preparation. The analysis of SSRs, effector genes, and mtDNA was insufficient to reveal the population structure of *P. infestans* isolates (Figure 1 and Supplementary Table S1).

Sequence variations in the *P. infestans* genome might contribute to its wide host range. In this study, while two isolates in the EGY2 cluster were members of 06_3928A from England [27], isolates in JPN1, JPN2, and EGY1 clusters were not represented in the major cluster comprising ten isolates analyzed in previous studies (Figure 1). Therefore, it is important to identify and classify the isolates correctly for successful plant disease management. Additionally, further investigation of effector genes is needed to explain the virulence profile of *P. infestans* isolates. Our results suggest that gene-based analyses are insufficient for isolate identification, and genotyping the isolates using genome-wide analyses, such as ddRAD-Seq and WGRS, is more powerful for the classification of new *P. infestans* isolates together with the previously reported isolates.

The number of isolates with a sequenced genome is limited; however, genome-wide sequence data of different isolates would provide more detailed phylogenetic information, which would help control isolates that could soon become endemic. Breeding late blight resistant tomato and potato cultivars is a promising strategy to overcome this disease [33, 34]. Characterization of *P. infestans* will enhance the monitoring of aggressive isolates propagated via infected plant materials from different geographic regions. Surveillance systems such as Asiablight, Euroblight, and USAblight have played important roles in monitoring the late blight disease [35]; however, similar programs are unavailable for other regions such as Africa. The plants were in fact completely damaged once infected by new isolates [36]. Overall, a short-term combat protocol based on prediction, diagnostics, and therapeutics is needed for sustainable farming and food production, and a long-term approach based on genetics, genomics, and plant breeding is needed for the effective control of late blight.

## Materials and Methods

### P. infestans isolates and DNA extraction

A total of 62 isolates were sampled from the leaves, stems, and fruits of tomato and potato plants growing in 16 geographical locations in Egypt during two growing seasons (2014– 2015 and 2015–2016) (Supplementary Table S1). Additionally, 18 isolates were obtained from the NIAS Genebank, Japan (http://www.gene.affrc.go.jp); these isolates were collected from the leaves and fruits of tomato plants and leaves of potato plants growing in three prefectures, Hokkaido, Ibaraki, and Shizuoka, from 1987–2013 (Supplementary Table S1). *P. infestans* isolates were extracted from the plants as described previously [33]. The DNA of isolates was extracted using the GeneJET Plant Genomic DNA Purification Mini Kit (Thermo Fisher Scientific) or DNeasy Plant Mini Kit (Qiagen, Hilden, Germany), according to the manufacturer’s instructions.

### Mating type assay

Mycelium plugs of *P. infestans* isolates and standard isolates of A1 or A2 mating type were placed on rye agar medium in a Petri dish at a spacing of 3 cm. All samples were incubated at 18°C in the dark for 3 weeks. The interaction domain of oospores between two isolates was investigated under a microscope at 20× magnification. Isolates that produced oospores (sexual spores) of the A2 mating type were identified as A1, while isolates that produced oospores of the A1 mating type were identified as A2 [37].

### Analysis of mtDNA and effector genes

The mtDNA of isolates was amplified from the genomic DNA using primers listed in Supplementary Table S2. The thermal cycling conditions used for PCR were based on previous studies [38]; [39]. Additionally, the presence/absence of five effector genes, including *AVR1, AVR2, AVR2-like, AVR3a*, and *AVR4*, in *P. infestans* isolates was tested in the current populations (Supplementary Table S2). The PCR products were separated by electrophoresis on 2% agarose gels and visualized using a UV gel documentation system (Biocraft, Co. Japan) after staining with ethidium bromide.

### SSR marker analysis

Sixteen SSR markers [40-42] were used to genotype *P. infestans* isolates using sequence-specific primers (Supplementary Table S2). DNA amplification by PCR and capillary electrophoresis using ABI3730 DNA analyzer (Applied Biosystems) were performed as described previously [24]. GeneMarker (SoftGenetics) was used to score the genotype. A phylogenetic tree was constructed based on the SSR data set using graphical genotypes (GGT) 2.0 software [43].

### Analysis of ddRAD-Seq data

Genomic DNA of isolates was double-digested with *Pst*I and *Msp*I restriction enzymes and used to construct ddRAD-Seq libraries. The libraries were sequenced using MiSeq (Illumina) in paired-end 251 bp mode, as described previously [44]; [33].

Data processing was performed as described previously [44] [33]. Low-quality sequences and adapters (Supplementary Table S2 and Supplementary Table S3) were removed from the raw sequence reads using PRINSEQ (version 0.20.4; [45] and fastx_clipper (FASTX Toolkit version 0.0.13; http://hannonlab.cshl.edu/fastx_toolkit), respectively. The cleaned reads were mapped onto the reference genome sequence of *P. infestans* strain T30-4 (ASM14294v1; accession number AATU00000000; [21] using Bowtie 2 (version 2.2.3; [46]. To obtain a variant call format (VCF) file containing SNP information, sequence alignment/map format (SAM) files were converted to binary sequence alignment/map format (BAM) files and subjected to SNP calling using the mpileup command of SAMtools (version 0.1.19; [47] and the view command of BCFtools [47]. High-confidence biallelic SNPs were selected using VCFtools (version 0.1.12b; [48], based on the following criteria: 1) ≥10× coverage in each isolate, 2) >999 SNP quality value, 3) ≥0.1 minor allele frequency, and 4) <0.5 missing data rate.

### Analysis of WGRS data

Sequencing libraries with an insert size of 600 bp were prepared for 17 isolates of *P. infestans*, as described previously [49] [33], and sequenced using Illumina NextSeq 500 in paired-end 151 bp mode. Data processing was conducted as described above for ddRAD-Seq data, with the exception of SNP filtering criteria, which were as follows: 1) ≥5× coverage in each isolate, 2) >999 SNP quality value, 3) ≥0.1 minor allele frequency, and 4) <0.2 missing data rate. Annotations of SNP effects on gene functions were predicted using SnpEff (version 4.2) [50]. CNVs were detected using CNV-seq (version 0.2.7) [51].

### Clustering analysis

Dendrograms were constructed using the unweighted pair-group method with average linkage (UPGMA). Additionally, PCA was performed to determine the relationship among samples using the TASSEL software [52], with the number of components limited to four.

## Accession numbers

Double digest restriction-site associated DNA sequencing (ddRAD-Seq) and whole genome resequencing (WGRS) data are available at the DDBJ Sequence Read Archive database under the accession number DRA007610.

## Acknowledgments

We thank the NIAS Genebank, Japan, for providing the *P. infestans* isolates. We also thank S. Sasamoto, C. Mimani, H. Tsuruoka, and A. Obara at the Kazusa DNA Research Institute for technical assistance. This work was supported by The Ministry of Higher Education and Scientific Research (MHESR), Egypt (Grant Number: 2013/2014-547), and the Kazusa DNA Research Institute Foundation, Japan.

## Supporting information captions

**Table S1.** *Phytophthora infestans* isolates used in this study.

**Table S2.** List of primer sequences used in this study.

**Table S3.** Number of ddRAS-Seq reads and rate of mapping against the reference genome sequence of *P. infestans*.

**Table S4.** Number of WGRS reads and rate of mapping against the reference genome sequence of *P. infestans*.

**Table S5.** Numbers of annotated SNPs and indels classified using the SnpEff program.

